# Development and application of a metabolomic tool to assess exposure of an estuarine amphipod to pollutants in the environment

**DOI:** 10.1101/2020.06.02.128983

**Authors:** Miina Yanagihara, Fumiyuki Nakajima, Tomohiro Tobino

**Affiliations:** Department of Urban Engineering, The University of Tokyo, 7-3-1 Hongo, Bunkyo-ku, Tokyo, Japan; Environmental Science Center, The University of Tokyo, 7-3-1 Hongo, Bunkyo-ku, Tokyo, Japan

**Keywords:** benthic organism, metabolomics, sediment toxicity, toxicity identification evaluation

## Abstract

Identifying substances in sediment that cause adverse effects on benthic organisms has been implemented as an effective source-control strategy. However, the identification of such substances is difficult due to the complicated interactions between liquid and solid phases and organisms. Metabolomic approaches have been utilized to assess the effects of toxicants on various organisms; however, the relationships between the toxicants and metabolomic profiles have not been generalized, and it makes impractical to identify major toxicants from metabolomic information. In this study, we used partial least squares discriminant analysis (PLS-DA) to investigate these relationships. The objective of this study was to construct PLS-DA models using the metabolomic profiles of the benthic amphipod *Grandidierella japonica* to assess the exposure effects of target toxicants and to demonstrate the utility of these models to assess the effects of chemicals in environmental samples. The PLS-DA models were constructed from the metabolomic profiles of *G. japonica* to discriminate the exposure of *G. japonica* to chromium, nickel, copper, zinc, cadmium, fluoranthene, nicotine, and osmotic stress. The models had high predictive power for the presumptive exposure to each chemical and were able to detect exposures in mixed chemical samples. These results suggest that the metabolomic responses can provide important information for the assessment of chemical effects on organisms. We applied the models to the metabolomic profiles of *G. japonica* exposed to river sediment and road dust, and the results demonstrated the applicability of the models. The control groups were not classified as exposure groups, and no samples were presumed to belong to any exposure group. These results suggest that the target chemicals were not toxic in the samples and conditions we investigated. This study demonstrates a method to assess the relationships between chemical exposure and metabolomic responses. To the best our knowledge, this is the first study to apply metabolomics for identification of toxicants.

## INTRODUCTION

Sediment contamination is widespread, and the effects of substances in the sediment on aquatic ecosystems are concerning. Preserving benthic ecosystems is important for maintaining biodiversity in aquatic habitats. Sediment quality guidelines have been proposed^1^, and it is possible to assess sediment toxicity, but the identification of major toxicants in the sediment is important to take a source-control management strategy. Urban runoff and road dust, which are major sources of pollutants in urban areas, has been investigated to assess the toxicity. Previous studies have reported the toxic effects of urban runoff on aquatic organisms^2,3^. They reported that aquatic ecosystems have been of concern, with major toxicants including heavy metals,^4,5^ organic substances,^5^ or polycyclic aromatic hydrocarbons (PAHs).^6^ Similarly, the major toxicants in road dust, a constituent of urban runoff, might consist of various chemicals including heavy metals,^7^ hydrophobic compounds,^8^ and organic compounds.^9^

Two approaches have been proposed to identify the major toxicants in the sediment - toxicity identification evaluation (TIE)^10^ and effect directed analysis (EDA).^11^ TIE for whole sediments has identified organic compounds as the major toxicants, and differences have been reported between the major toxicants of whole sediments and interstitial water.^12^ The TIE approach uses adsorbents to separate the samples into several fractions, and the EDA approach is based on fractionation, both of which can complicate the assessment of the bioavailability of substances in the sediment. Therefore, another approach based on biological information is required to identify the major toxicants. A biomarker is a biological response that can be used to assess the status of an organism. For example, metallothionein is a well-known protein with a high affinity for heavy metals, and it has been used to detect heavy metal exposure by analyzing changes in the protein levels. This is also true for aquatic invertebrates – changes in metallothionein levels have been used to assess heavy metal exposure.^13^ However, there are limitations to this approach. The increases in protein levels are not specific to a heavy metal, which makes it difficult to identify the toxicant. Thus, more biomarkers are required to observe complicated biological responses.

Metabolomics has been used in ecotoxicology to observe the subcellular responses of test organisms. This approach comprehensively analyzes low-molecular substances in organisms and enables the elucidation of reactions in their bodies.^14^ In most cases, the test organisms are exposed to target toxicants or any physiological stressors and the metabolomes from the body are measured by mass spectrometry or nuclear magnetic resonance spectroscopy.^15^ These measurements allow insights into the status of the organism, even when the target compounds or genome information is unknown. The relationships between each toxicant and the metabolomic profile have been investigated by metabolomics; however, the specific relationships remain unclear. Many studies have reported that metabolites change due to several factors, but the metabolites have been inconsistent. For example, when *Daphnia magna* was exposed to copper at similar concentrations in two studies, one reported an increase in N-acetylspermidine and threonine,^16^ whereas the other reported an increase in glycerophosphocholine.^17^ Similarly, when *D. magna* was exposed to different concentrations of fluoranthene, different metabolites showed significant changes.^18^ When the bivalve *Corbicula fluminea* was exposed to a mixture of zinc (Zn) and cadmium (Cd), the results were complicated, and the effect of mixing was unclear.^19^ The inconsistencies in the metabolites may be due to correlations among the metabolomes, for which univariate analyses (i.e., t-test) are ill-suited. Thus, a new methodology is required to elucidate the relationships between the metabolomic profiles and the exposure substances to identify the major toxicants.

This study aimed to develop a new tool for screening sediment toxicity in the environment, with two objectives. The first objective was to build models for the estimation of toxicants affecting estuarine amphipods, using metabolomic profiles of the amphipod. The target toxicants include substances from road dust^9,20,21^ such as heavy metals (chromium [Cr], nickel [Ni], copper [Cu], Zn, and Cd) and organic chemicals (fluoranthene and nicotine). Osmotic stress was also included because changes in salinity can affect the toxicity of chemicals in road dust.^22^ The models were assessed with training and test data sets and investigated for their applicability to a single chemical and a mixture of chemicals. The second objective was to assess the applicability of the models to the toxicants in sediment samples collected from an urban area (i.e., road dust and river sediment samples). The estuarine amphipod, *Grandidierella japonica*, was used as a test species in this study, because it has been used in acute and chronic sediment toxicity tests.^23,24^ This species is indigenous to Japan but has recently been documented in the Mediterranean Sea.^25^

## MATERIALS AND METHODS

### Test organisms and sediment samples

*Grandidierella japonica* was collected from Nekozane River in Chiba, Japan in 2009 as previously described,^26^ and the culture was used for exposure tests. The amphipods have been cultured with artificial seawater (30‰) and sediment collected from the Komatsugawa tidal flats.^24^ The sediment was sieved by a stainless sieve (mesh size: 2.0 mm) and autoclaved at 121°C before stored in a refrigerator (4 °C). They are fed with grounded TetraMin^®^ three times per week, and the culture has been maintained followed by the same method mentioned in the previous research.^22^

River sediment and road dust samples were collected for the following exposure testing. The information on sediment and road dust is summarized in Table 1. Two river sediment samples were collected from two tidal areas; the bottom of the canal (ES1 sample) and river (ES2 sample) in Tokyo. Two road samples (RD1, RD3 samples) were collected from the metropolitan expressway by road sweeping vehicles as previously reported.^9,23^ The concentrations of heavy metals and organic pollutants are summarized in Table S1.

**Table 1.**
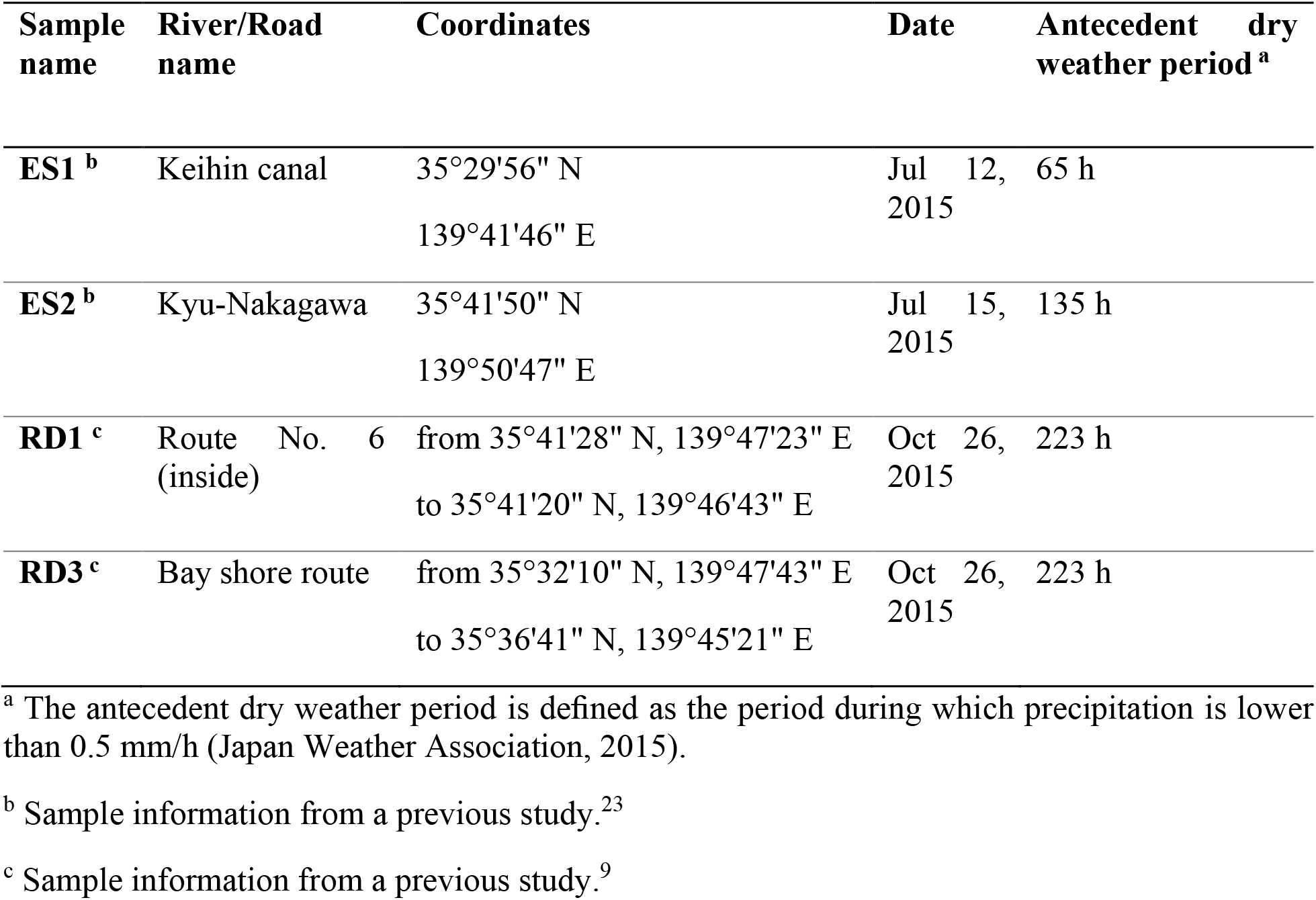
Sediment sample information

### Four-day exposure tests to obtain metabolomic data for the discriminant models

Toxicity tests with Cr, Ni, Cu, Zn, Cd, fluoranthene (Flu), nicotine (Nic), and osmotic stress (Sal) were conducted to obtain the metabolomic information for the discriminant models. Six stock solutions were prepared with deionized water by dissolving a chemical out of potassium dichromate (K_2_Cr_2_O_4_), nickel(II) chloride hexahydrate (NiCl_2_ · 6H_2_O), copper sulfate pentahydrate (CuSO_4_ · 5H_2_O), zinc sulfate heptahydrate (ZnSO_4_ · 7H_2_O), cadmium nitrate tetrahydrate (Cd(NO_3_)_2_·4H_2_O), and nicotine. A stock solution of fluoranthene was prepared using dimethyl sulfoxide (DMSO) because of the low solubility to water, and DMSO was used as the solvent control for the fluoranthene exposure test. Test water was made by adding the stock solutions into artificial seawater (30‰) so that the nominal concentration was 50% lethal concentration (LC50) (High-dose group) or 10% of LC50 (Low-dose group). The LC50 was obtained from previous reports^9,27,28^ and additional experiments (unpublished data). The conditions were assigned according to the chemical name (e.g., Cu, Zn) and concentration (e.g., High-dose or Low-dose). In addition to the 14 exposure groups, we also included a “Mix” exposure group, created by mixing the Cu, Zn, and Cd stocks at the LC50 level (High-dose) and at 10% of LC50 (Low-dose). Finally, a salinity group (Sal) was prepared to examine the effects of changing salinity concentrations at 5‰ (Low-dose) or 45‰ (High-dose). We based the exposure concentrations on LC50 to obtain metabolomic information under comparable situations; this has also been used in transcriptomic studies.^29^

Four-day exposure testing with the prepared test water was conducted by modifying a standardized method.^24^ For each chemical, we set up three beakers for the Control group and three to five beakers for the exposure group. The conditions for the exposure tests are summarized in Table 2. Each beaker contained 120 mL of test water and 30 g of quartz sand, and the aeration was not conducted in the tests. After one day of incubation (25 ± 1 °C, 16 h light/8 h dark cycle), 10 *G. japonica* juveniles were added to each beaker. The juvenile amphipods were collected using a 710 and a 500 μm mesh net^9^ and were not fed during the exposure tests. After four days, the surviving juveniles were collected in vials (three to five individuals per vial). Detailed information on the mortality is shown in Table 2. The vials were flash-frozen using liquid nitrogen and stored at −80 °C until metabolome extraction.

**Table 2.**
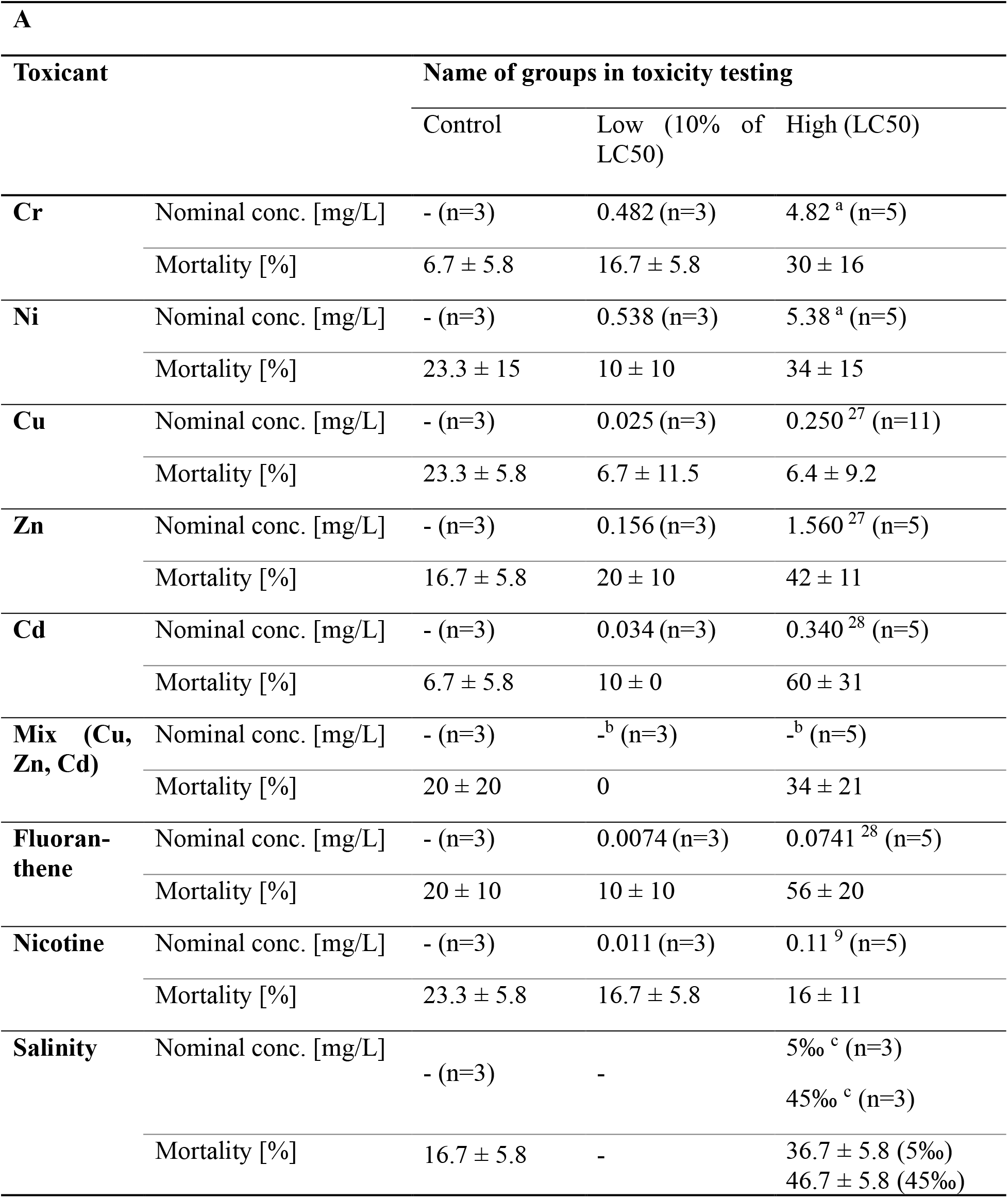
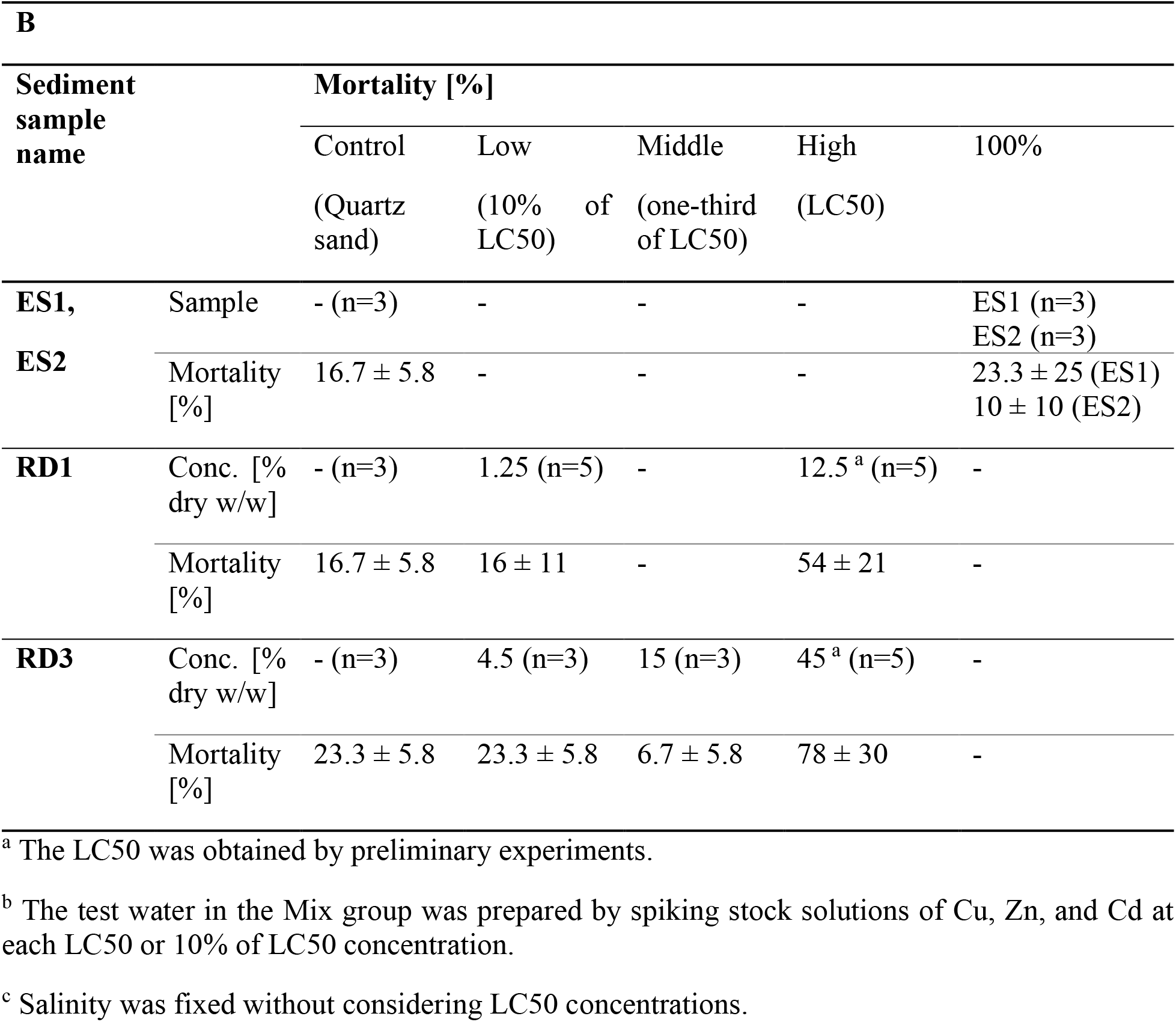
The detailed information on the four-day exposure tests. (A: Tests for model construction, B: Tests for model application; the nominal concentration and the number of replicates in each test are shown with the mortality of the test (mean ± S.D.).)

### Four-day exposure to environmental samples to assess the applicability of the discriminant models

*Grandidierella japonica* was exposed to river sediment (ES1 and ES2) and road dust samples (RD1 and RD3) as in the exposure tests explained above. Each beaker contained 30 g of test sediment and 120 mL of artificial seawater. The overlying water was aerated during these tests. The test sediment was composed of quartz sand (Control groups), river sediment (ES1 or ES2), or a mixture of road dust and quartz sand (RD1 or RD3). The river sediment was used without dilution because a previous study (Hiki et al., 2019) reported the mortality after 10-day exposure testing was lower than 50%. The road dust was highly toxic, so the samples were mixed with quartz sand for concentrations at LC50 (High-dose), 33% of LC50 (Middle-dose), and 10% of LC50 (Low-dose). The detailed condition of the tests and the results of mortality are shown in Table 2. Four-day exposure tests were conducted, and the surviving individuals were used for metabolomic analysis.

### Extraction and analysis of the metabolomes from the amphipod samples

Metabolomes of *G. japonica* were extracted using water, methanol, and chloroform, as described in a study.^26^ The polar fraction was used for metabolomic analysis, and the samples were analyzed using the Exactive Orbitrap mass spectrometer (Thermo Fisher Scientific, Waltham, MA, USA) following a previously outline method.^30^ The samples were analyzed by flow-injection and the flow rate of mobile phase was 200 μL/min. The *m/z* range was set to 100 – 1,000 to detect low-molecular weight substances. The molecular weights and signal intensities of the metabolomes were analyzed using SIEVE software (version 2.2 SP2; Thermo Fisher Scientific). The intensity was pre-processed as previously described^30^ and used as explanatory variables in multivariate analysis. Briefly, the fold change was obtained by comparing the normalized intensity between the control and exposure groups and auto-scaled prior to the data analysis.

### Multivariate analysis of the metabolomic data

Principal component analysis (PCA) and partial least squares discriminant analysis (PLS-DA) were conducted using the pre-processed values from the High-dose exposure groups. We focused on data from the High-dose groups to observe major changes compared to the Control groups so that we could build a screening tool. First, we used PCA to illustrate the characterization of metabolomic responses of *G. japonica* exposed to different chemicals. Second, we constructed eight PLS-DA models to discriminate a given chemical from the other chemicals. The PLS-DA method can be used to supervise the correlation between variables, even when the set of explanatory variables is noisy and large.^32^ The explanatory variables were the pre-processed signal values of the metabolites, and the response values were labeled 0 or 1. The metabolomic data was separated into two groups; training data set (High-dose groups) and test data set (Control and Low-dose groups). We conducted another analysis under different condition; the data from the Low-dose exposure group was used as training data set and High-dose groups and Control groups were regarded as test data set, for reference (details provided in Supplementary Material). For example, the Cu model was built to separate the Cu exposure group (High-dose) from the other exposure groups (High-dose). In this example, the Cu and Mix exposure groups were labeled as 1, and the other groups were labeled 0. Similarly, we constructed models to discriminate exposure to Cr, Ni, Cu, Zn, Cd, fluoranthene, nicotine, and salinity with training data set. The number of latent variables was selected according to Wold’s R criterion^33^, the Q^2^ values were calculated for all models to assess their predictive power,^34^ and the variable importance in projection (VIP) values were obtained for all compounds within each model.^31^ The VIP values were used to filter the important variables that contributed to the regression. The eight models were rebuilt with the variables selected by the VIP values, and the predictive power was checked by Q^2^ values. Finally, the test data set was used to assess the classification performance of the models. Chemical names and formulas of the detected compounds were taken from the Kyoto Encyclopedia of Genes and Genomes (KEGG) database^35^ based on the monoisotopic mass (mass error < 10 ppm).

## RESULTS

### Comparison of the metabolomic profiles of G. japonica between different factors

We conducted PCA using the pre-processed intensity values of 7,474 metabolites, and the results showed differences in metabolomic responses due to different factors. The score plots, shown in Figure 1, indicate the major effects of osmotic stress because the Sal group (5‰) was explained by PC1 (Figure 1a). Figure 1b shows the relationships between PC2 and the fluoranthene and nicotine groups. The differences were smaller among the heavy metal exposure groups (Cr, Ni, Cu, Zn, and Cd). The effects of osmotic stress, fluoranthene, and nicotine exposure were more pronounced probably because of their different modes of action. The PCA results indicate that multivariate analyses could be used to extract more information from the metabolomic data.

**Figure 1.**
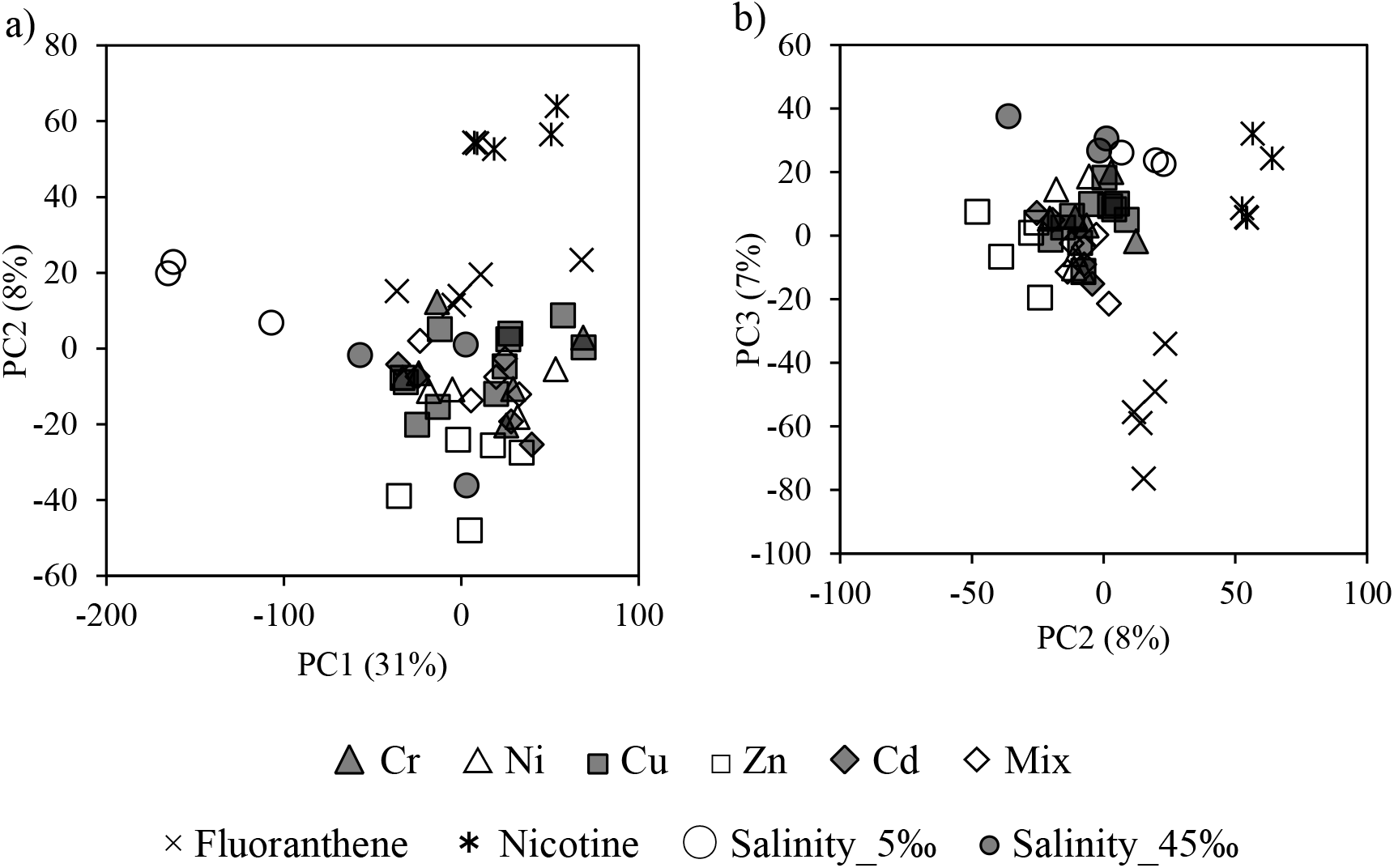
PCA score plots. (a: Score plot with PC1 and PC2, b: Score plot along with PC2 and PC3)

### PLS-DA models

The characteristics of the metabolomic profiles were used to discriminate individual exposures by PLS-DA. We developed eight PLS-DA models using 7,040 metabolites as explanatory variables in the High-dose exposure groups. The Q^2^ value for the Cr and Ni models was lower than 0.5, suggesting low predictive power. However, the Q^2^ values for the other models were higher than 0.5 and ranged between 0.66 and 0.97. PLS-DA distinguishes which variables contribute to the correlation between explanatory and response variables by calculating VIP values. We extracted the metabolites with higher importance by selecting metabolites with VIP > 1.5. The number of selected variables was 344, 691, 228, 464, 520, 830, 577, and 339 for the Cr, Ni, Cu, Zn, Cd, Flu, Nic, and Sal models, respectively. We conducted PLS-DA with the selected variables and rebuilt the eight models, resulting in Q^2^ > 0.5 in all models (Table 3). The rebuilt models were used as subsequent discriminant models owing to their high predictive power.^34^ The VIP-selected variables differed among the models (Figure 2). The metabolites with the five highest VIP values are shown in Table 4.

**Table 3.**
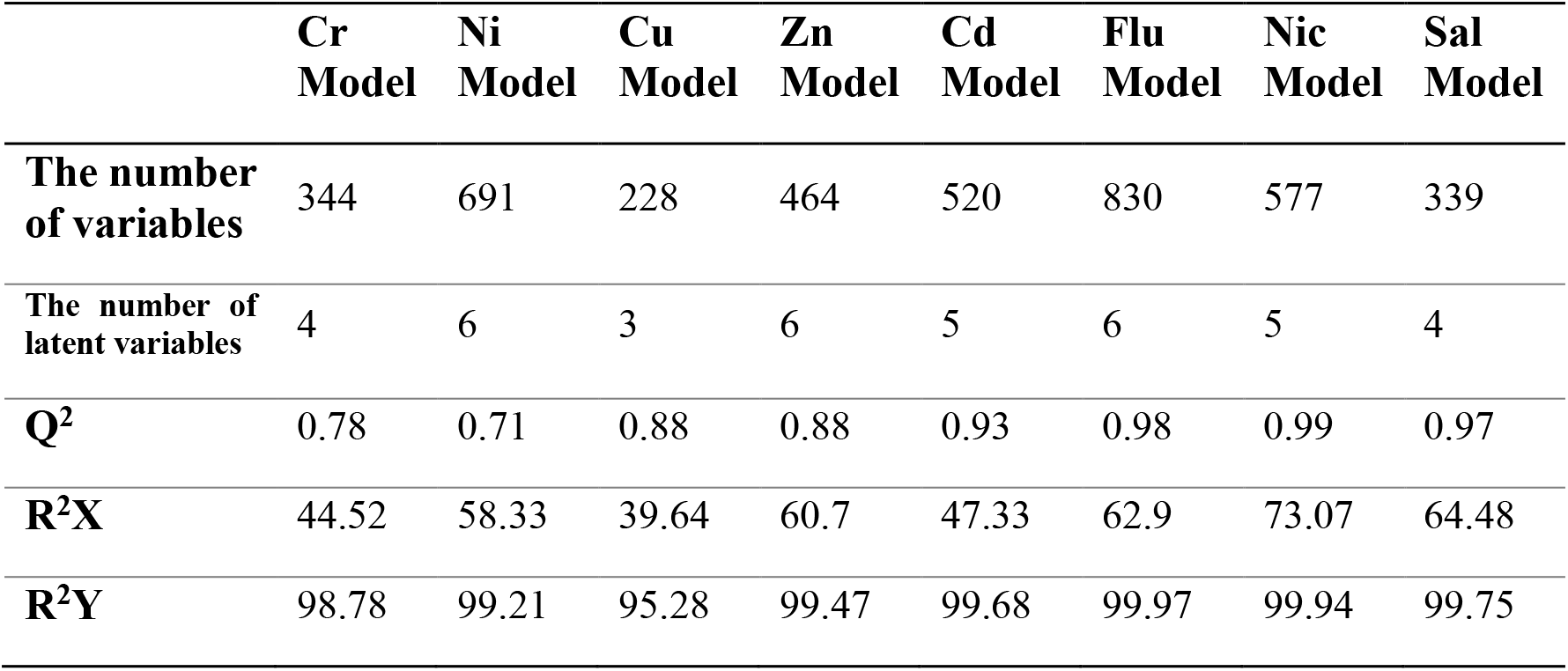
Properties of the PLS-DA models

|  | **Cr**<br>**Model** | **Ni**<br>**Model** | **Cu**<br>**Model** | **Zn**<br>**Model** | **Cd**<br>**Model** | **Flu**<br>**Model** | **Nic**<br>**Model** | **Sal**<br>**Model** |
| --- | --- | --- | --- | --- | --- | --- | --- | --- |
| **The number**<br>**of variables** | 344 | 691 | 228 | 464 | 520 | 830 | 577 | 339 |
| **The number of**<br>**latent variables** | 4 | 6 | 3 | 6 | 5 | 6 | 5 | 4 |
| **Q<sup>2</sup>** | 0.78 | 0.71 | 0.88 | 0.88 | 0.93 | 0.98 | 0.99 | 0.97 |
| **R<sup>2</sup>X** | 44.52 | 58.33 | 39.64 | 60.7 | 47.33 | 62.9 | 73.07 | 64.48 |
| **R<sup>2</sup>Y** | 98.78 | 99.21 | 95.28 | 99.47 | 99.68 | 99.97 | 99.94 | 99.75 |

**Figure 2.**
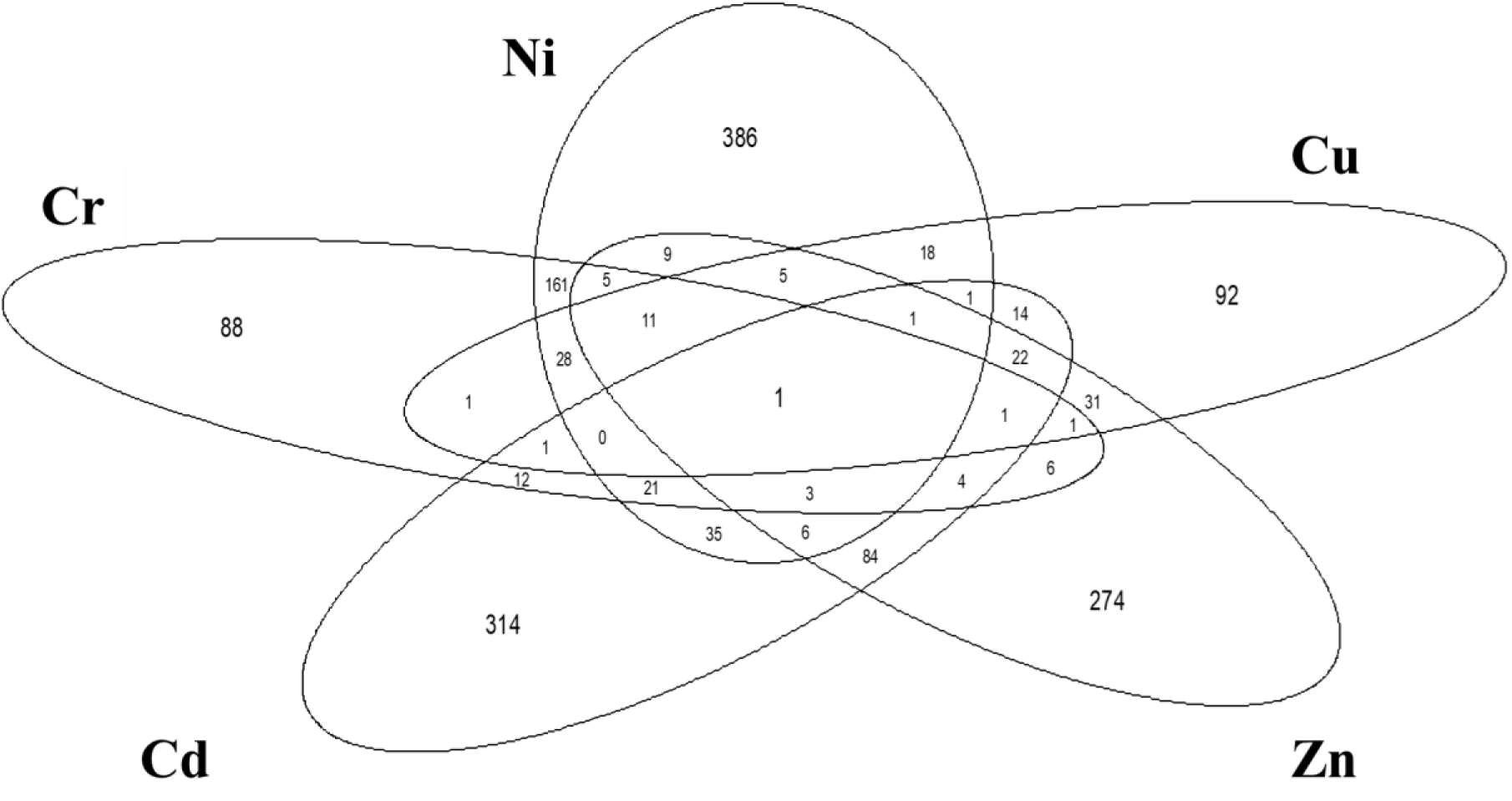
Commonality of the metabolites with VIP higher than 1.5 in the models to detect heavy metal exposure

**Table 4.**
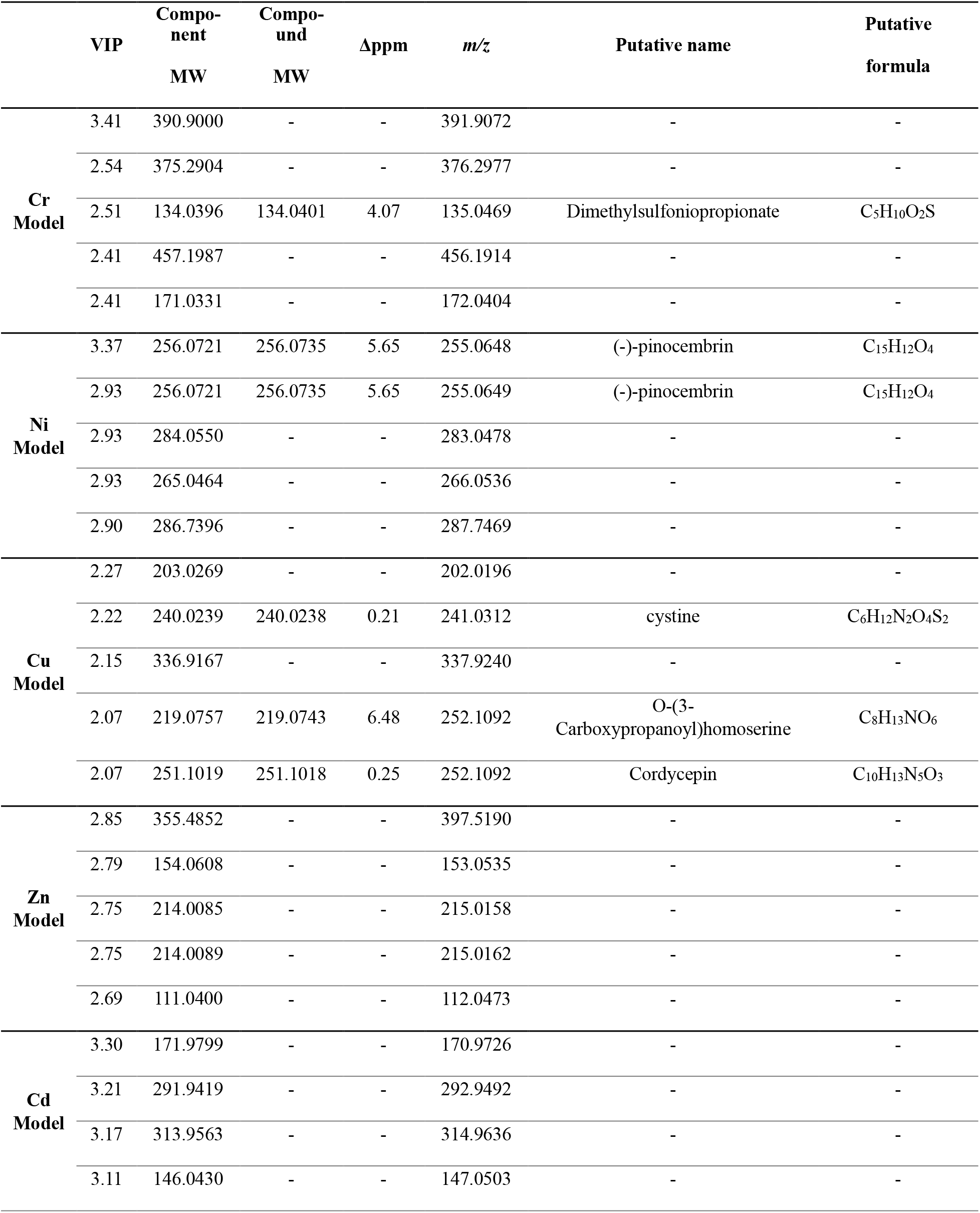
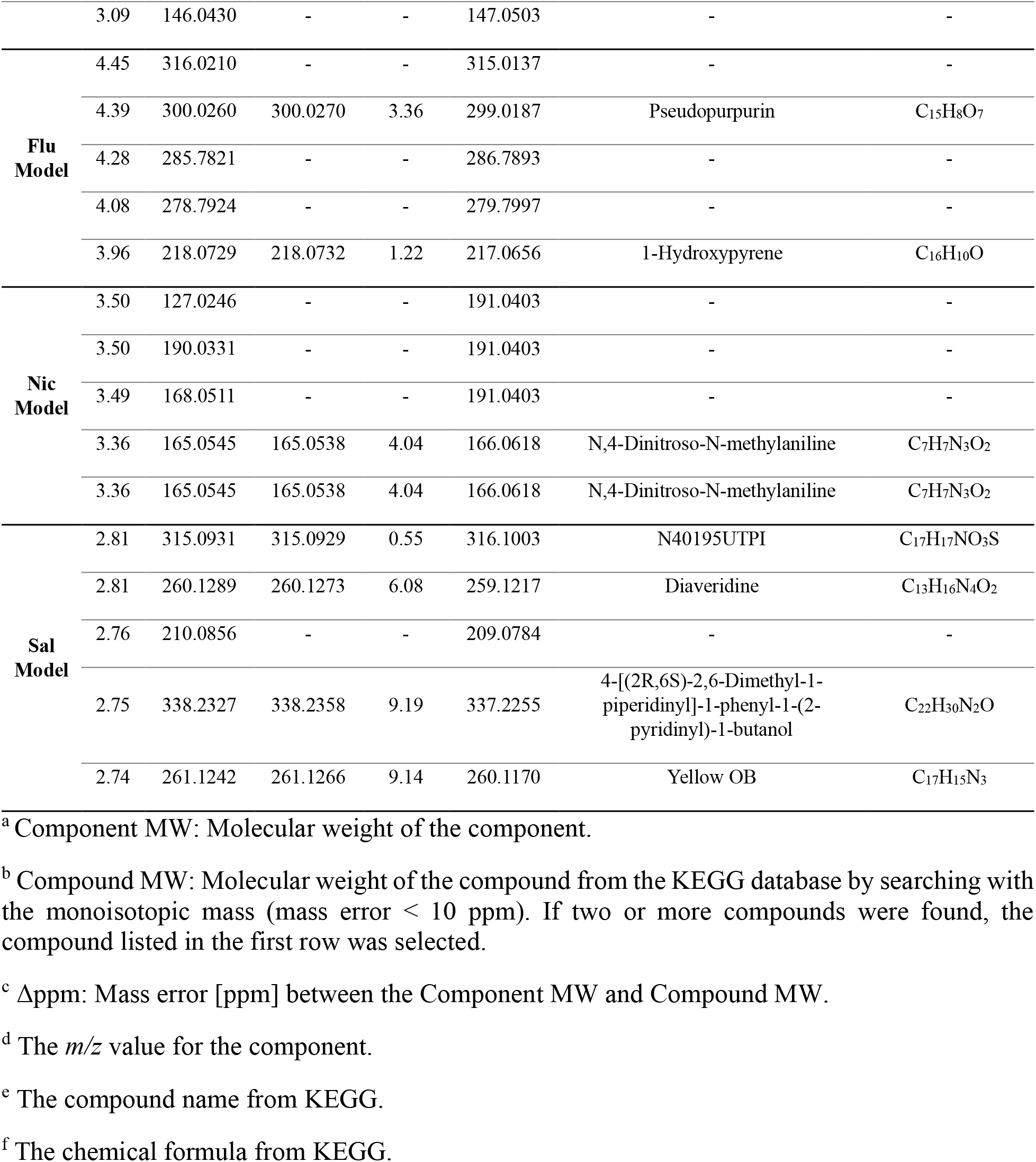
Significant metabolites from each model that contributed to the discrimination of the target toxicant. The metabolites with the five highest VIP values are shown in the table.

### Assessment of capacity of discrimination for each model

We calculated the response values of the rebuilt models using the training data set (High-dose groups) and test data set (Control and Low-dose groups). The mean of 1 and 0, 0.5, was used as threshold to classify the response values in Figure 3. Samples in the High-dose group of the target chemical are expected to show response values > 0.5, whereas samples in the High-dose group of the other chemicals are expected to show response values < 0.5. As shown in Figure 3, the training data set was classified as trained, compared to 0.5. The response values of most of the samples were regressed to around 1 or 0 in all models. For example, in the Cr model, the response values of the Cr exposure group (High-dose) were close to 1, and the values of the other exposure groups (High-dose) were close to 0. However, the Cu model showed a large variance in the values, which may be related to low mortality in exposure testing. The Mix exposure group was classified into the target chemical exposure groups in the Cu, Zn, and Cd models. Therefore, these models demonstrated the ability to discriminate chemicals in the mixture.

**Figure 3.**
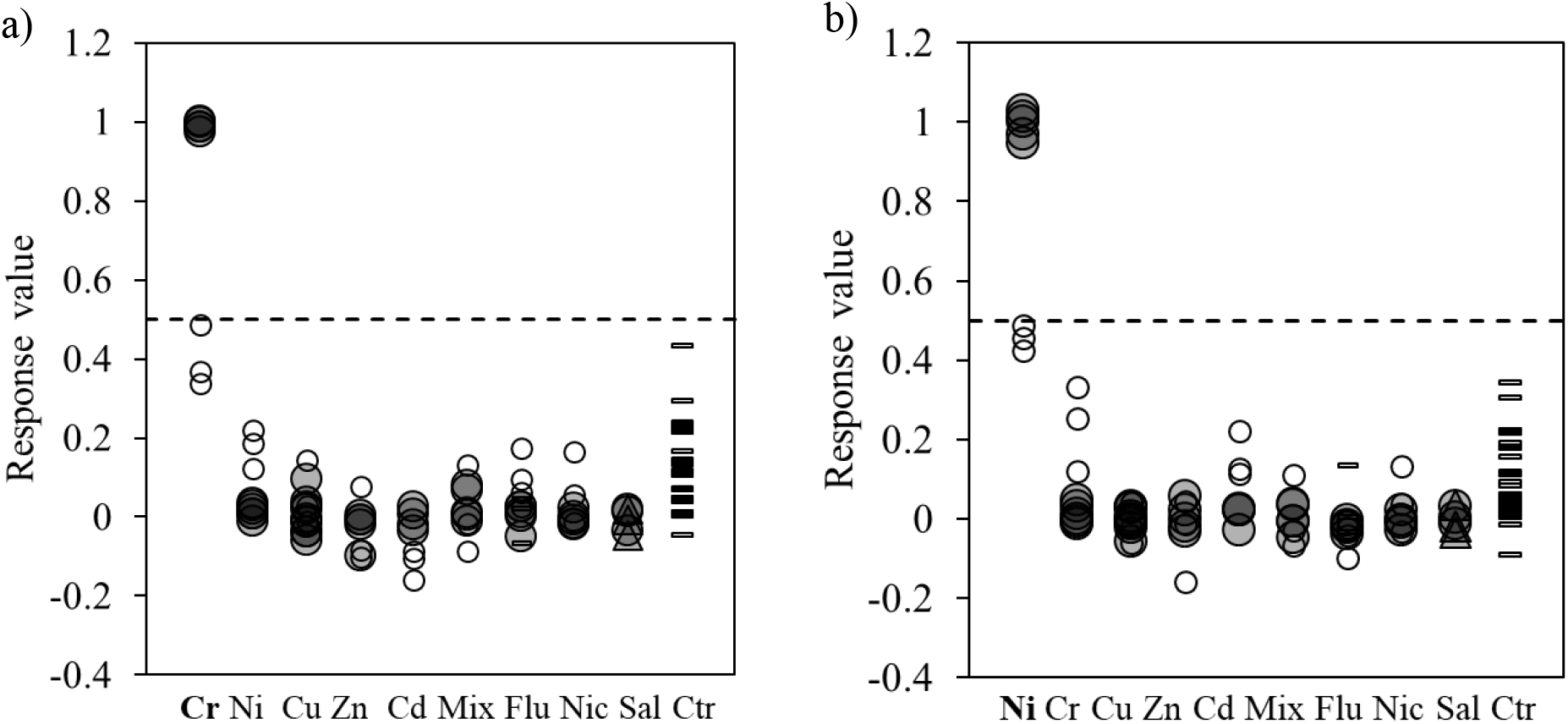
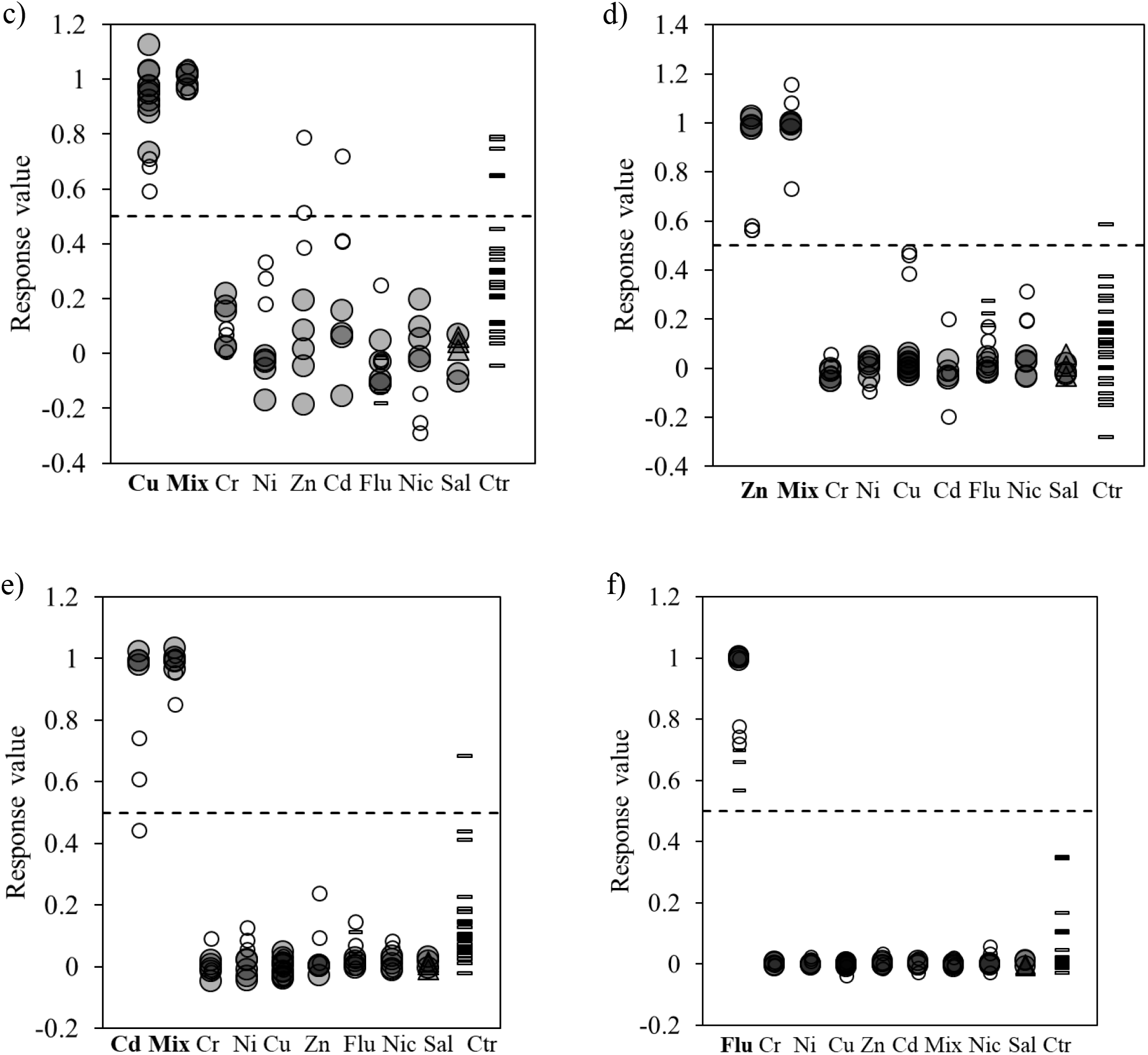
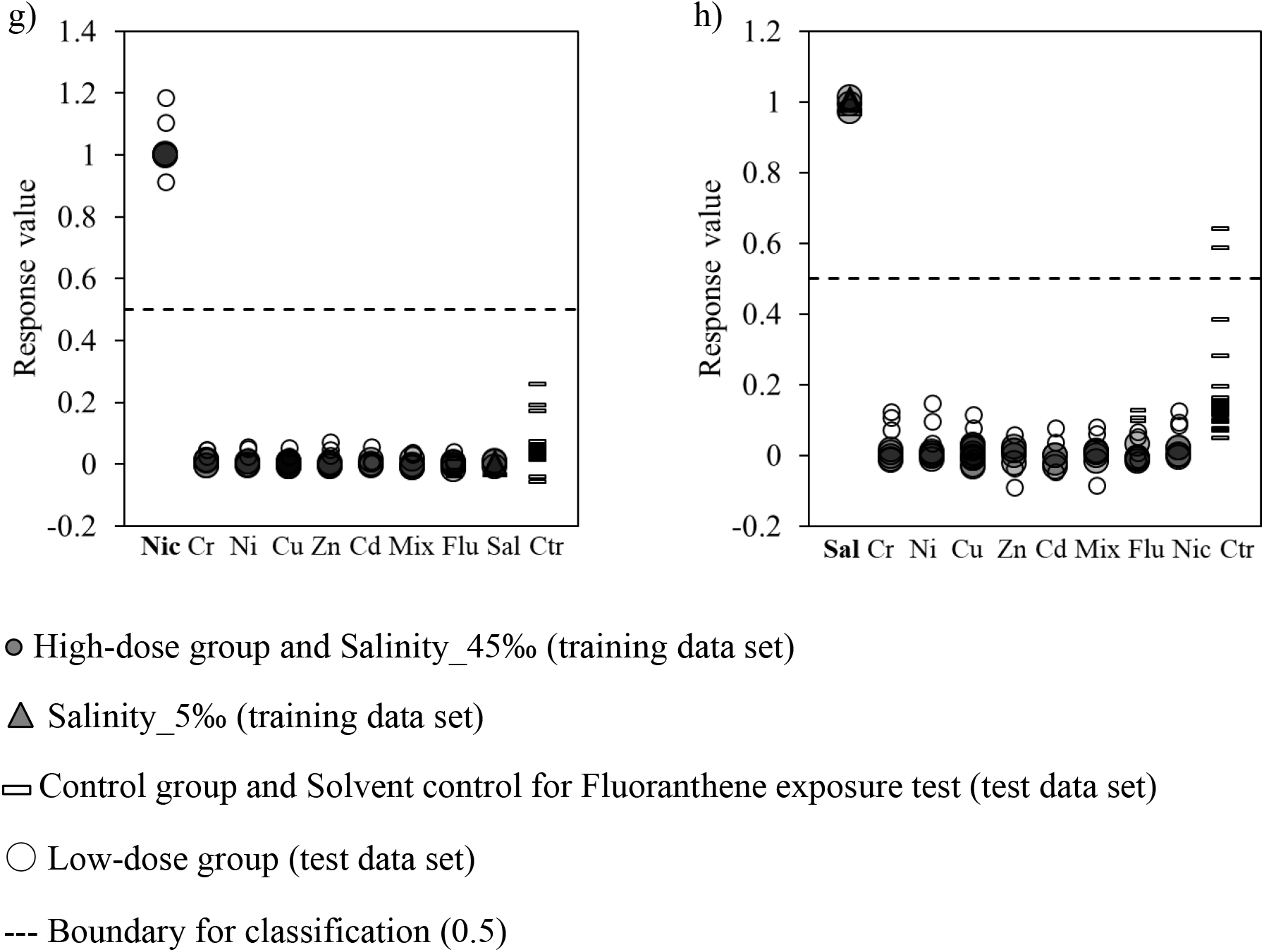
The response values of the discriminant models (a: Cr, b: Ni, c: Cu, d: Zn, e: Cd, f: Flu, g: Nic, h: Sal)

The models were validated with the test data set, and the classification capability of the models were assessed. The response values in the Control groups were < 0.5 in 95% of the samples, on average. However, the response values in the Cu model tended to vary more than the other models, which was consistent with the result of the training data set. The values of the samples in the Low-dose exposure groups varied among the models. In the Cu, Zn, Cd, Flu, and Nic models, some samples in the Low-dose exposure group of the target chemical had values > 0.5. The results of the test data set suggest that Control groups can be classified as non-exposure groups in almost all models. The classification performance of the Low-dose groups was dependent on models; the Zn, Cd, Flu, and Nic models could detect exposure effects at low-dose exposure levels. There may be relationships between the metabolomic responses and exposure concentrations, and further experimental investigations are needed to clarify the relationships. Another analysis that used Low-dose groups as training data set showed low performance of classification (Fig. S1 in Supplementary Material). It suggests that extrapolation of the effects at lower levels to the effects at higher levels is not suitable in this experiment.

### Application of the models to assessing exposure of chemicals in sediment samples

We conducted another metabolomic analysis to apply the models built above and confirmed the applicability of the models to the assessment of the environmental samples. The metabolomic profiles obtained from *G. japonica* exposed to either of ES1, ES2, RD1, or RD3 were input to eight models. The response values in each model are shown in Figure 4. First, all Control samples showed lower response values than 0.5 in all models except the Cu Model. Given that the accuracy of the Cu Model was low, this result indicated that the results of the Control were reproducible, and the discriminant models based on the information of metabolomic profiles were able to be applied to assessment of a new data set. The response values of most of the samples showed lower values than 0.5, and it indicated that those samples were not exposed to any target chemicals. The response values of several samples in Control, ES2, and RD1 groups in the Cu Model showed a higher value than 0.5. However, the Cu Model had an error in discrimination of Control case in Figure 3, and we could not judge the classification.

**Figure 4.**
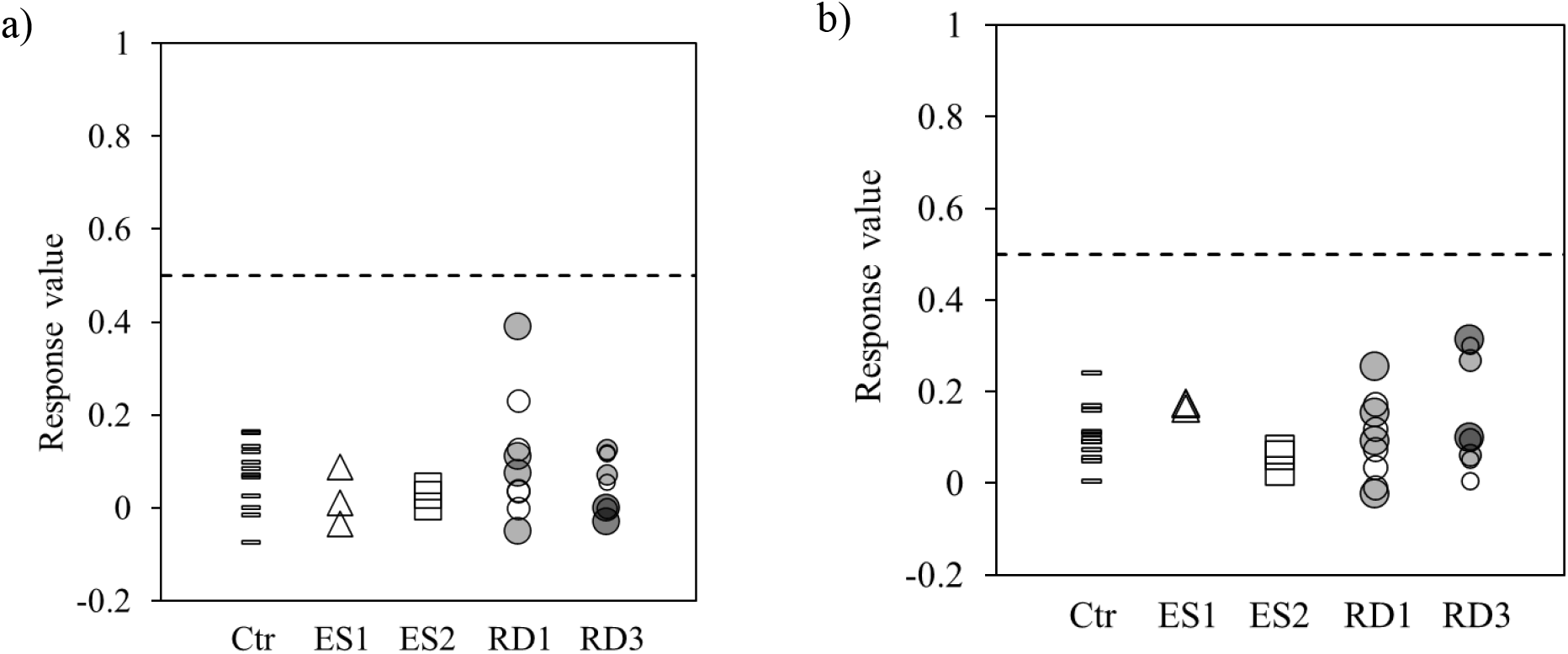
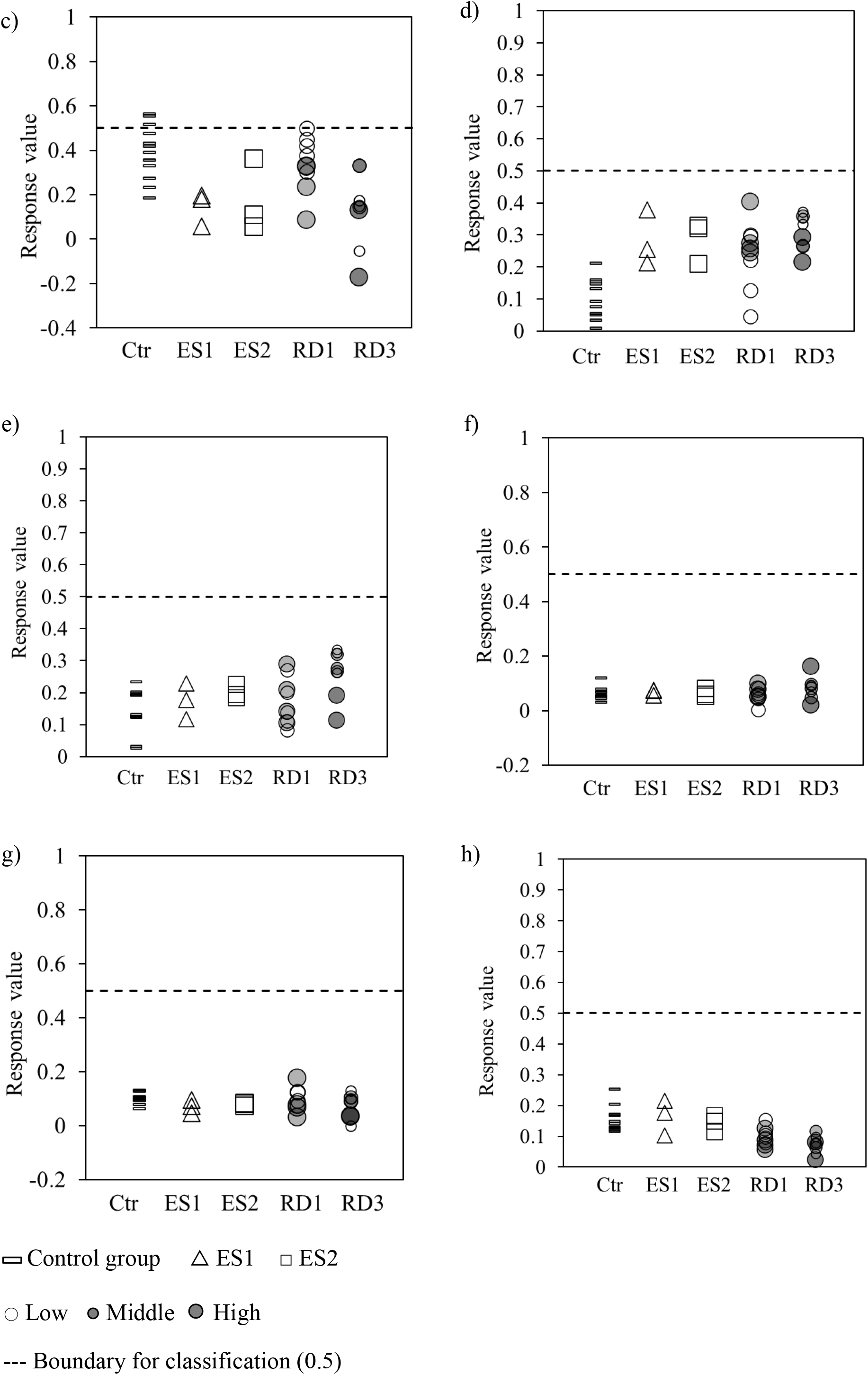
The response values of the discriminant models based on the metabolomic profiles of *G. japonica* exposed to sediment samples. (a: Cr, b: Ni, c: Cu, d: Zn, e: Cd, f: Flu, g: Nic, h: Sal)

## Discussion

### Multivariate analysis for elucidating the metabolomic responses

We compared the metabolomic profiles by PCA, and the score plots (Figure 1) showed that differential chemical exposure might cause different metabolomic responses in *G. japonica*. We applied those characteristics to interpret the information for the exposure to substances by PLS-DA. As a result, we obtained eight models that discriminated the exposure groups (Table 3). Several previous studies have reported variable changes in different metabolites due to chemical exposure. However, investigations of the specific relationships between the metabolites and chemicals have been challenging. For example, two studies analyzed the metabolites of *D. magna*^16,17^ and reported variations in different metabolites after copper exposure. However, these results were from univariate analyses, while other studies suggest that the metabolites may be related.^36^ This may explain the inconsistencies in previous studies. The PCA and PLS-DA results indicate that multivariate analyses are better suited to extract important information from large data sets of metabolomic profiles.

### Annotation of the metabolites with high VIP values

PLS-DA provides a VIP value for each metabolite to rank the contribution of the explanatory variables. We calculated VIP for each metabolite in each model and built new models with metabolites that had VIP > 1.5. This resulted in higher Q^2^ values, suggesting that the selection of variables by VIP is an effective method of noise reduction. The commonality of the metabolites was assessed by a Venn diagram (Figure 2), but nearly all of the compounds differed among the models. This suggests that the metabolites used to discriminate exposure to different chemicals tend to be different. As discussed above, previous studies have shown that many metabolites change after exposure to chemicals. Therefore, we employed VIP-based selection to rank the relative importance of the metabolites.

The putative names of some metabolites with VIP > 1.5 were identified from the KEGG database. We identified adenosine in the Cr, Ni, Cd, and Flu models, arachidonic acid in the Cr, Ni, Cu, and Zn models, and leucine in the Ni, Zn, Cd, Flu, Nic and Sal models. Adenosine is a nucleotide that increases in *Corbicula fluminea* exposed to a mixture of Cd and Zn^19^ and in *Mytilus galloprovincialis* exposed to Ni or chlorpyrifos.^37^ However, adenosine decreases in *M. galloprovincialis* exposed to the mixture of Ni and chlorpyrifos. These results suggest that adenosine responds to many chemicals and may lack specificity. Arachidonic acid, an unsaturated fatty acid, decreases in *D. magna* exposed to Cd, dinitrophenol, and fenvalerate and increases in *D. magna* exposed to proporanolol.^38^ Leucine is an amino acid that widely varies, but the specific relationships are unclear. For example, leucine increases in *D. magna* exposed to copper^17,39^ and lithium^17^ and varies with levels of starvation, higher temperature, and bacterial infection in *Haliotis rufescens*.^40^ Similar results have been reported for *Mytilus edulis* exposed to atrazine and starvation stress,^41^ *M. galloprovincialis* exposed to chlorpyrifos and nickel,^37^ *M. galloprovincialis* exposed to mercury and PAHs,^42^ and *Ruditapes philippinarum* exposed to arsenic.^43^ We successfully selected the variables, but most of the metabolites with the five highest VIP values (Table 4) were not identified. Identification of the metabolites requires further investigation to achieve a better understanding of the metabolic pathways.

### Relationships between the exposure concentrations and the predictive power of the models

The PLS-DA models were tested by training and test data sets; the training data set was trained successfully (Figure 3). The proposed models could detect the exposure effects of Cr, Ni, and Salinity at LC50 levels and the effects of Zn, Cd, fluoranthene, and nicotine at 10 % of LC50 levels. The models were built using training data set (High-dose groups) and a tool to screen severe effects was obtained. Another analysis that used Low-dose groups as training data set was not successful (Figure S1 in Supplementary Material), probably because metabolomic responses that detected in Low-dose groups were not strongly related to each target toxicant.

The control groups in the test data set showed response values < 0.5 in all models, except the Cu model. This suggests that most of the models can be utilized to classify the control groups based on the metabolomic profiles. The Low-dose groups in the test data set showed variable response values. The exposure concentrations of Cu, Zn, Cd, fluoranthene, and nicotine (Low-dose) were lower than the no observed effect concentration (NOEC) and the lowest observed effect concentration (LOEC) for *G. japonica,^9,27,44^* suggesting that the metabolomic responses were sensitive enough to be observed at concentrations lower than NOEC. This may be an advantage of this approach - it allows exposure detection at a wide range of concentrations. Investigations of the relationships between the exposure concentrations and metabolomic responses are promising avenues to inform more useful discriminant models.

### Applicability of the discriminant models to assess environmental samples

The second objective of this study was to apply the models to the assessment of environmental samples. We used the models to assess chemical exposure in river sediment (ES1, ES2) and road dust (RD1, RD3) samples. The response values of the control samples were < 0.5 in most models (Figure 4), suggesting that the models apply to the new samples. Compared with the chemical property of the samples,^9,23^ the result of the models seemed to be inconsistent with the presence of the target chemicals in the sediment samples. The models were built based on the metabolomic responses, and they should include the consideration of bioavailability. Therefore, the results of the models suggest that the target chemicals in ES1, ES2, RD1, and RD3 samples may not contribute to the toxic effects at LC50 levels. Moreover, regarding Zn, Cd, fluoranthene, and nicotine, the exposure effects might be smaller than effects at 10% of LC50 levels, according to the comparison between the response values in Figures 3 and 4.

To the best of our knowledge, the present study is the first to propose and apply discriminant models based on the metabolomic responses of macroinvertebrates. Several transcriptomic studies have discussed discriminant methodology, but none have demonstrated applicability to a mixture of chemicals in environmental samples or the detection of individual target chemicals.^29,45^ Our discriminant models using metabolomic profiles successfully identified toxicants when *G. japonica* was exposed to a single chemical or a mixture and showed the applicability to screen the toxic effects in the sediment samples collected from the environment.

Here, we proposed the models that can be used to screen severe effects of the target toxicants, and further work needs to be performed to elucidate two things; the range of the concentrations and exposure route that are detected by the models. All of the sediment samples contained the target toxicants^9,21^, but the results suggest that the exposure effects of the target chemicals in the sediment samples were lower than the effects at LC50. Also, a previous study^9^ suggests major effects of nicotine in a road dust sample, though the models in this study did not show the same result. More research on these points would help us to establish more effective models.

## CONCLUSIONS

We conducted exposure testing and metabolomic analyses using *G. japonica* and investigated the relationships between the chemical information and metabolomic profiles. We utilized the metabolomic responses of *G. japonica* to predict the effects of exposure by PLS-DA, generating eight discriminant models for exposure predictions to Cr, Ni, Cu, Zn, Cd, fluoranthene, nicotine, and osmotic stress. The models using fewer metabolites with VIP values >1.5 had higher predictive power. The models were able to discriminate a single chemical in a mixture, and the boundaries for classification were defined based on the training data set. Subsequently, the models were used to assess the effects of the target chemicals in river sediment and road dust samples. The models successfully classified the control samples into non-exposure groups, indicating that the models can be used to detect toxic effects. The response variables of the exposure groups suggested no major effects of the target factors. To the best of our knowledge, this metabolomic approach has never been used in ecotoxicology - we provide the first report of a discriminant analysis method based on metabolomic profiles.

## AUTHOR INFORMATION

### Author Contributions

MY, FN, and TT designed this study. MY conducted the experiments including exposure testing and metabolomic analysis and wrote the initial draft of the manuscript. FN and TT contributed to interpreting the data and critically reviewed the manuscript. All authors approved the final version of the manuscript.

## ACKNOWLEDGMENT

This work was supported by the Steel Foundation for Environmental Protection Technology (SEPT), Japan.

