## Supplementary Material for "Development and application of a metabolomic tool to assess exposure of an estuarine amphipod to pollutants in the environment"

This document includes the results of the information on the concentrations of the target toxicants in the sediment samples and results of the models that were built with Low-dose data set.

*Detailed information on the target toxicants in the tested sediment samples*

The concentrations of the target toxicants in the sediment samples have been reported by previous studies.[9,23] The information is summarized in Table S1.

Table S1 Concentrations of target toxicants in sediment samples. (A single asterisk and double asterisks indicate the value is greater than ERL and ERM, respectively.)

| | **Concentrations in sediment samples [mg/kg-dry]** | | | | | | |
|---|---|---|---|---|---|---|---|
| | Cr | Ni | Cu | Zn | Cd | Fluoranthene | Nicotine |
| **ES1** [a] | 29 | *39.7 | *127 | *407 | 1.1 | 0.22 | - |
| **ES2** [a] | *357 | **67.7 | *263 | **703 | *2.13 | *2.43 | - |
| **RD1** [b] | *163 | **96 | *150 | **1200 | 0.26 | 0.23 | 8.45 |
| **RD3** [b] | *123 | *37.3 | *133 | **783 | 0.34 | 0.08 | 0.76 |
| **ERL** [c] | 81 | 20.9 | 34 | 150 | 1.2 | 0.6 | - |
| **ERM** [c] | 370 | 51.6 | 270 | 410 | 9.6 | 5.1 | - |

[a] Sample information from a previous study.[23]

[b] Sample information from a previous study.[9]

[c] ERL: Effect range low, ERM: Effect range medium.[1]

*PLS-DA models with Low-dose data set*

The High-dose data set obtained from *G. japonica* was used to build PLS-DA models in the manuscript. Here, Low-dose data was used as training data set and the response values of the other data were assessed. Figure S1 shows the result of the response values of the test and training data set. We have also applied the models to predict toxicity of the environmental sediment and road dust. The result of the application of the models to the environmental samples is shown in Figure S2. As shown in Figure S1, the training of Low-dose data set was conducted successfully. However, the response values of the High-dose data set varied and the classification tended to fail. It suggests that the extrapolation of the metabolomic responses may be difficult with Low-dose data set, compared with the result of the main manuscript. Also, as detailed in Figure S2, the response values of all models were lower than 0.5 when the models were applied to assessment of the target toxicants. Great care must be taken when discussing the results of Figure S2 because the models showed low performance of classification. The further investigation on the dose-response relationships at metabolites-levels is essential toward the development of useful assessment tool.

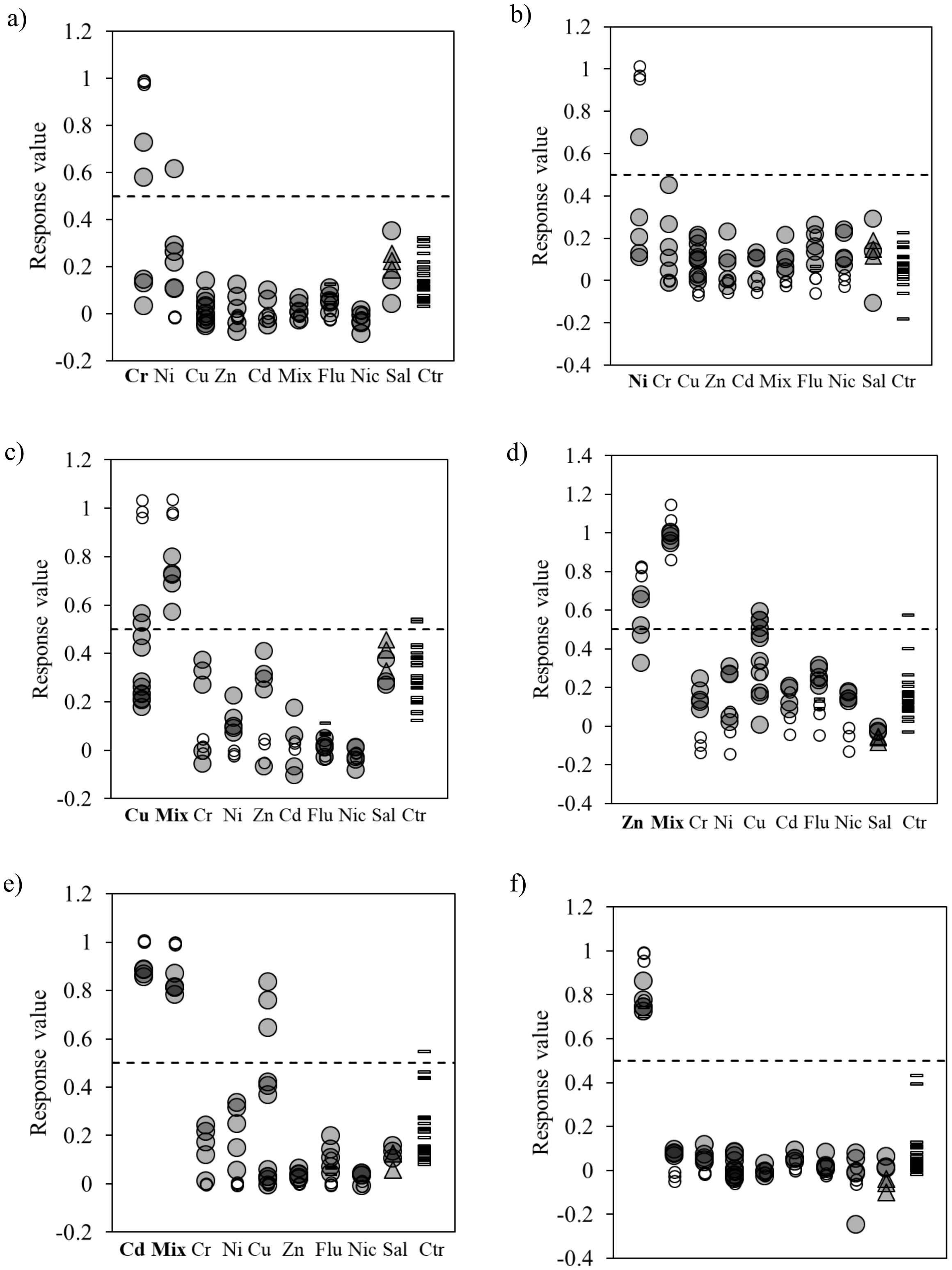
a)
Response value
Cr Ni Cu Zn Cd Mix Flu Nic Sal Ctr
b)
Response value
Ni Cr Cu Zn Cd Mix Flu Nic Sal Ctr
c)
Response value
Cu Mix Cr Ni Zn Cd Flu Nic Sal Ctr
d)
Response value
Zn Mix Cr Ni Cu Cd Flu Nic Sal Ctr
e)
Response value
Cd Mix Cr Ni Cu Zn Flu Nic Sal Ctr
f)
Response value
Flu Cr Ni Cu Zn Cd Mix Nic Sal Ctr

g)

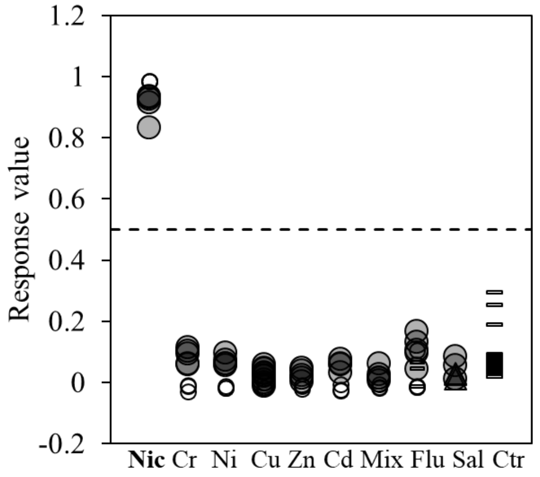


● High-dose group and Salinity_45‰ (training data set)

▲ Salinity_5‰ (training data set)

▭ Control group and Solvent control for Fluoranthene exposure test (test data set)

○ Low-dose group (test data set)

--- Boundary for classification (0.5)

**Figure S1.** The response values of the discriminant models based on the metabolomic profiles of *G. japonica* exposed to sediment samples (a: Cr, b: Ni, c: Cu, d: Zn, e: Cd, f: Flu, g: Nic)

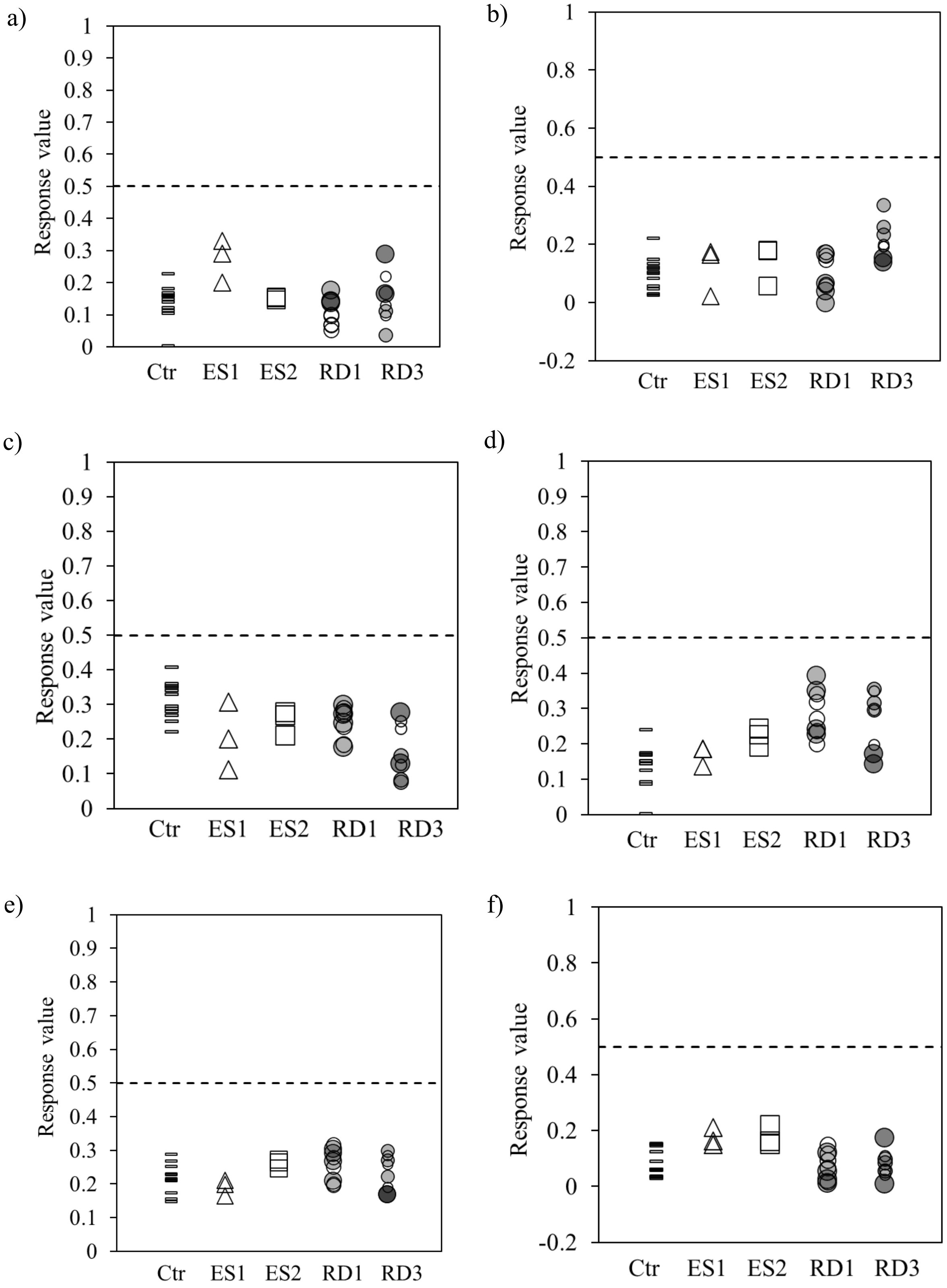
a)
1
0.9
0.8
0.7
0.6
0.5
0.4
0.3
0.2
0.1
0
Response value
Ctr
ES1
ES2
RD1
RD3
b)
1
0.8
0.6
0.4
0.2
0
-0.2
Response value
Ctr
ES1
ES2
RD1
RD3
c)
1
0.9
0.8
0.7
0.6
0.5
0.4
0.3
0.2
0.1
0
Response value
Ctr
ES1
ES2
RD1
RD3
d)
1
0.9
0.8
0.7
0.6
0.5
0.4
0.3
0.2
0.1
0
Response value
Ctr
ES1
ES2
RD1
RD3
e)
1
0.9
0.8
0.7
0.6
0.5
0.4
0.3
0.2
0.1
0
Response value
Ctr
ES1
ES2
RD1
RD3
f)
1
0.8
0.6
0.4
0.2
0
-0.2
Response value
Ctr
ES1
ES2
RD1
RD3

g)

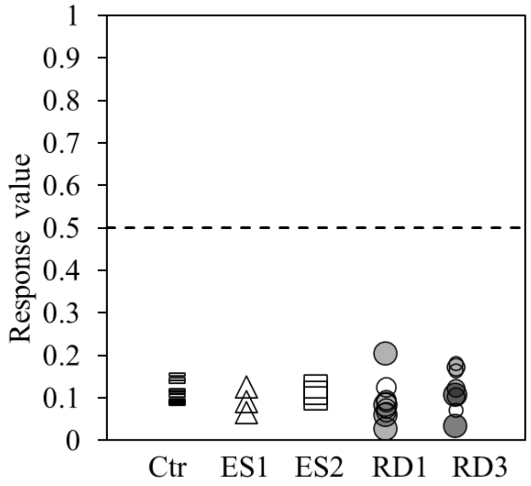


Control group △ ES1 □ ES2

○ Low ● Middle ● High

--- Boundary for classification (0.5)

**Figure S2.** The response values of the discriminant models based on the metabolomic profiles of *G. japonica* exposed to sediment samples. (a: Cr, b: Ni, c: Cu, d: Zn, e: Cd, f: Flu, g: Nic)
